# Pseudo-linear Summation explains Neural Geometry of Multi-finger Movements in Human Premotor Cortex

**DOI:** 10.1101/2023.10.11.561982

**Authors:** Nishal P. Shah, Donald Avansino, Foram Kamdar, Claire Nicolas, Anastasia Kapitonava, Carlos Vargas-Irwin, Leigh Hochberg, Chethan Pandarinath, Krishna Shenoy, Francis R Willett, Jaimie Henderson

## Abstract

How does the motor cortex combine simple movements (such as single finger flexion/extension) into complex movements (such hand gestures or playing piano)? Motor cortical activity was recorded using intracortical multi-electrode arrays in two people with tetraplegia as they attempted single, pairwise and higher order finger movements. Neural activity for simultaneous movements was largely aligned with linear summation of corresponding single finger movement activities, with two violations. First, the neural activity was normalized, preventing a large magnitude with an increasing number of moving fingers. Second, the neural tuning direction of weakly represented fingers (e.g. middle) changed significantly as a result of the movement of other fingers. These deviations from linearity resulted in non-linear methods outperforming linear methods for neural decoding. Overall, simultaneous finger movements are thus represented by the combination of individual finger movements by pseudo-linear summation.

## Introduction

Dexterous finger movements are fundamental to many everyday living activities, enabling tasks as simple as picking up an apple to as complex as playing a Chopin nocturne. Although the full range of hand movements requires more than 20 degrees of freedom, many complex hand movements can be more simply decomposed into a combination of flexion/extension movements of each finger joint (Rahman and Al-Jumaily 2013). The principles that govern how the neural representation of these individual movements is combined to produce complex, multi-finger movements remain largely unknown.

One hypothesis is that neural representation for a movement is equal to a simple combination of the neural representation of its parts (‘compositional coding’). The simplest compositional hypothesis for multiple finger movements is the linear summation of constituent single finger movements (Figure 1A). This hypothesis has been successful in explaining the motor cortical representation of homologous movements across limbs as the linear summation of a movement-specific and a limb-specific component (Willett et al. 2020; Guan et al. 2023).

**Figure 1.**
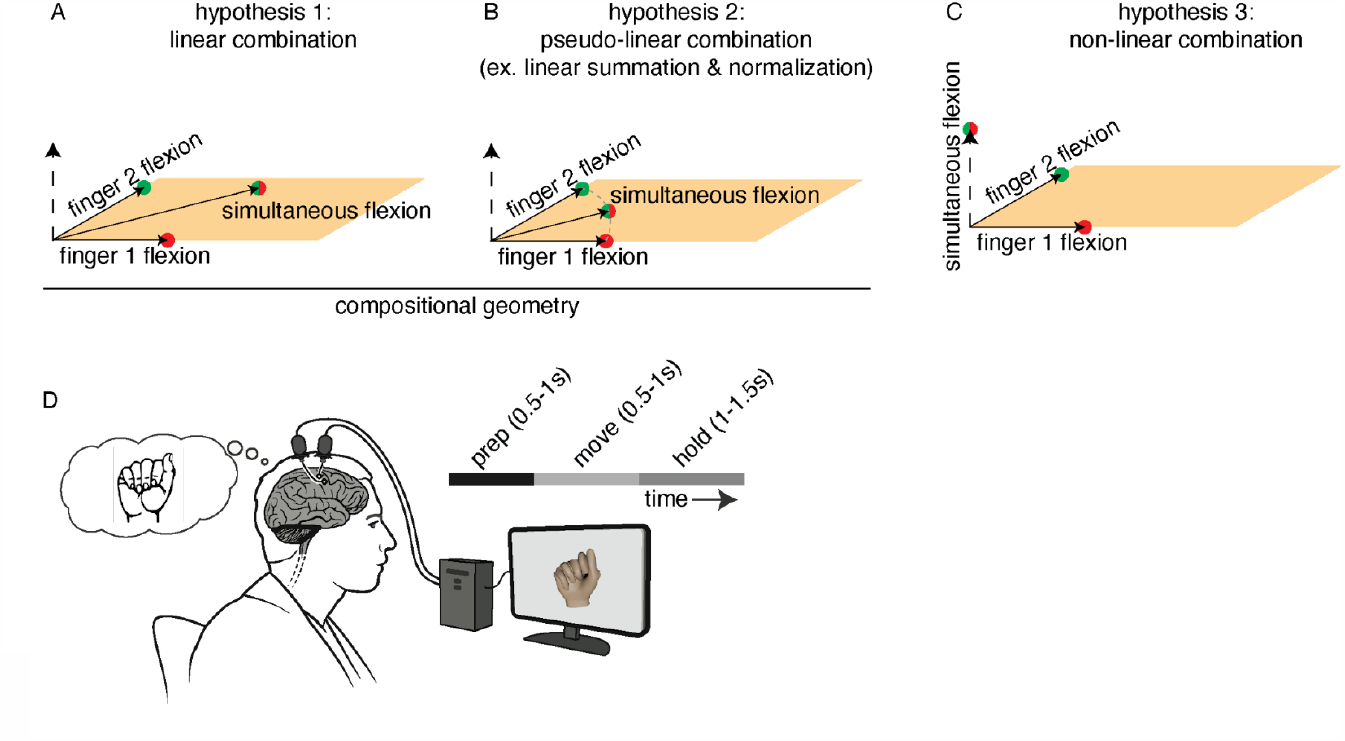
Hypotheses and research setting: (A, B, C) Hypotheses for the neural geometry of multiple finger movements in neural state space, where each axis corresponds to the activity of a neuron. (A, B) represent compositional coding, where the neural activity for multiple finger movements is equal to the activity for single finger movements combined by simple operations such as linearity summation (A) or pseudo-linear summation (B). (C) Neural activity for multiple finger movements is a non-linear combination of single finger movement activities, such that the resulting activity is largely orthogonal to the subspace spanned by single finger movements in the neural state space.(D) Research setup. Intracortical neural activity is recorded using two Utah arrays placed in the hand-knob area of the dominant precentral gyrus while the participant attempts finger movements cued using an animated hand in 3D.

An alternative hypothesis is that the relationship between the neural representation of multiple finger movements and single finger movements is nonlinear and not predictable in a simple way from the component parts. This hypothesis can be geometrically described in the neural state space, with the axes corresponding to the activity of neurons in the circuit. In this model, the neural activity representing single and multiple finger movements exists in orthogonal subspaces (Figure 1C), allowing a large number of movements to be distinctly represented along unique neural dimensions.

The compositional coding hypothesis in its simplest form predicts that the magnitude of neural activity increases with the number of moving fingers, which may be unrealistic in a biological neural circuit, in which firing rates of individual neurons reach a saturation limit. To address this potential issue, a more sophisticated compositional hypothesis adjusts the initial simple linear summation by adding magnitude normalization (‘pseudo-linear’ summation, Figure 1B). This revised compositional hypothesis limits representational capacity as a large number of movements are constrained to a low dimensional space spanned by single finger movements.

We investigated these hypotheses using intracortical multi-electrode array recordings from the human premotor cortex as research participants with paralysis attempted single, paired or higher order finger movements (Figure 1D). Our participants repeatedly attempted a wide variety of multi-finger movements designed to explore a broad parameter space including both natural gestures and arbitrary combinations of flexion and extension.

Our results were most consistent with the pseudo-linear compositional hypothesis in which the neural activity for multiple finger movements was aligned with the linear summation of constituent single finger movement activities, but the magnitude was lower compared to linear summation (consistent with normalization).

## Results

Intracortical neural recordings were obtained from two participants in the BrainGate2 pilot clinical trial (‘T5’ and ‘T11’) as they attempted single finger and multi-finger movements. Both T5 (69 years old) and T11 (36 years old) are right-handed men, with tetraplegia secondary to a high-level spinal cord injury. Both had two 96-channel intracortical Utah arrays placed in the ‘hand knob’ area of the left precentral gyrus. All research sessions focused on the fingers of the right hand.

### Neural geometry of single finger movements

We began by characterizing the neural representation of single finger movements. Participant T5 attempted movements of the right hand (contralateral to the arrays), with trials starting from a neutral rest position and ending in either attempted flexion or extension of a single finger. The movement for each trial was cued using a 3D animated hand and consisted of preparatory (1.5s) and move (1s) periods (Figure 1D). Movements were repeated across three research sessions resulting in a total of ∼1000 trials.

Neural activity on single electrodes, measured by the number of threshold crossing events, differed across the movement conditions, confirming tuning for single finger movements (Figure 2A). Both finger velocity and finger position were represented in the neural activity (Supplementary Figure 1). Neural population activity occupied a low-dimensional space, with five principal components (out of 576 ambient dimensions = concatenation of 192 channels of trial-averaged activity across three sessions) capturing > 95% of the variance (Figure 2B).

**Figure 2.**
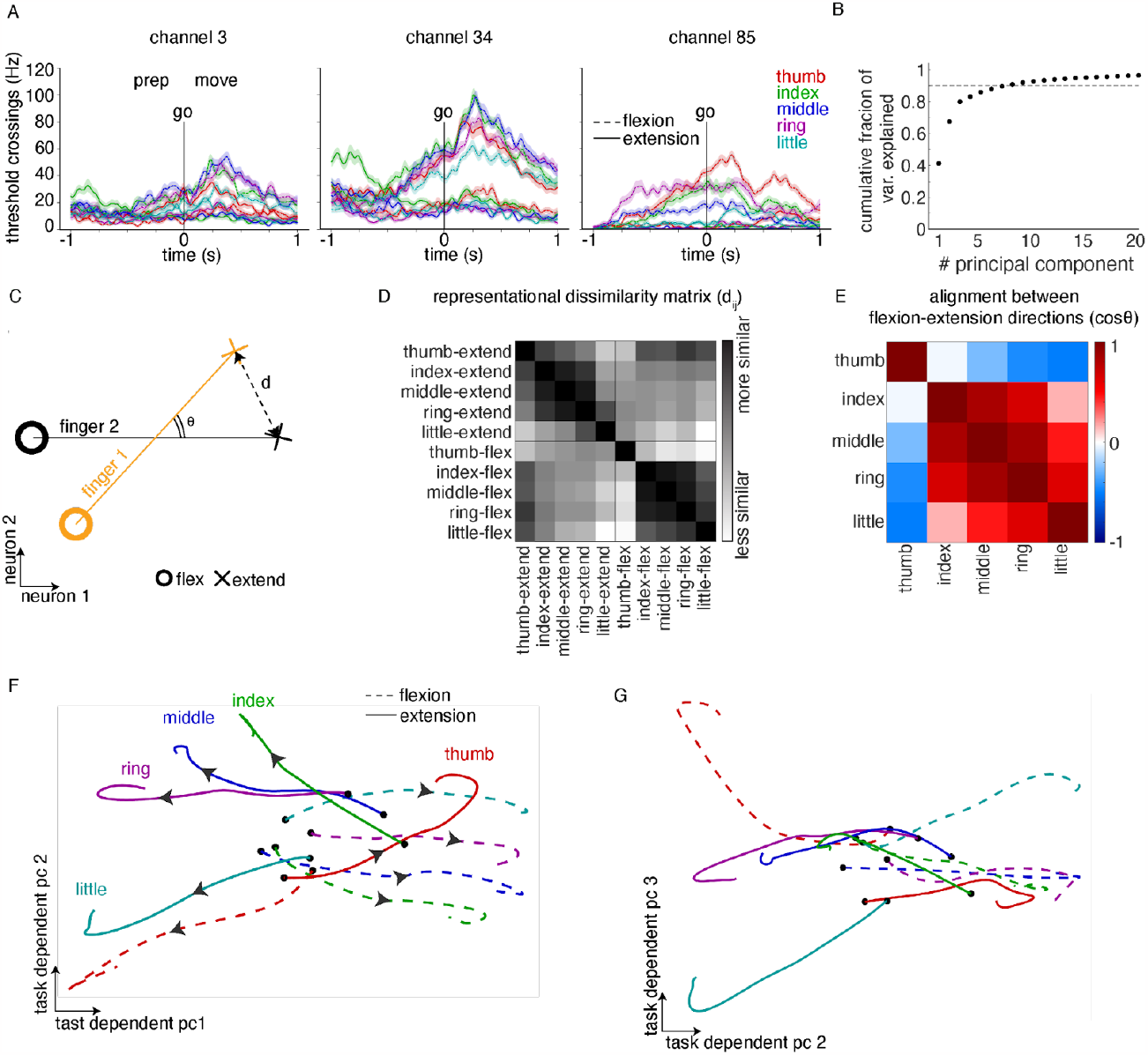
Neural geometry of single-finger movements. (A) Peristimulus time histograms of recorded neural activity from 3 example channels during attempted flexion and extension of individual fingers on the contralateral (right) hand (lines show the average for different conditions, shaded regions show standard error). (B) Cumulative fraction of variance explained by the principal components of the trial and time-averaged neural activity. The top five components captured > 90% of the variance (ambient dimensionality 576 resulting from concatenation of 192 channels across three sessions), suggesting a low dimensional representation. (C) Schematic describing two metrics for characterizing the geometry of different movements in neural state space, where each axis corresponds to the activity of a particular neuron. Representational dissimilarity matrix (RDM) is estimated by measuring Euclidean distance (d_ij_) between pairs of movements. Relative arrangement of the direction along which each finger modulates the activity in the neural state space is given by the cosine of angle (θ) between lines joining flexion-extension movements of a given finger. (D) RDM, averaged across three sessions. Black indicates higher similarity. (E) Correlation between modulation directions for pairs of fingers. Nearby fingers are correlated, except for the thumb. (F, G) Visualization of neural activity using cross validated Targeted Dimensionality Reduction (TDR) followed by PCA. TDR directions identified by regressing from the kinematic variables (five dimensions, one for each finger, +1 for flexion/ -1 for extension and 0 for rest) to trial & time-averaged neural activity in the last 0.4s of each trial. Colors indicate fingers, line style indicates different movements (flexion/extension) and black dots indicate the ‘go’ cue. Arrows indicate the evolution of neural activity during the course of the trial. Two views of the 3D geometry are shown.

The geometry of different finger positions in the neural state space was assessed using a Representational Dissimilarity matrix (RDM, (Kriegeskorte, Mur, and Bandettini 2008)), estimated by measuring the cross-validated Euclidean distance of the population activity between all pairs of movements (flexion/extension across fingers) after trial and time averaging (see Methods, Figure 2C). The distances between flexion and extension movements were large, resulting in a block diagonal structure (Figure 2D) and within each movement type, nearby fingers were more similar (Supplementary Figure 2). This structure is similar to the RDM of the kinematics and muscle activity during finger movements in able-bodied people (Ejaz, Hamada, and Diedrichsen 2015; Natraj et al. 2022; Guan et al. 2022).

In addition to the RDM, the cosine of the angle between the lines joining flexion and extension movements for each finger was used to measure the relative arrangement of the direction along which each finger modulates the activity in the neural state space (Figure 2E). While nearby fingers showed a greater correlation between their directions (as expected), the thumb was anti-correlated to the little finger, which was not expected based on previously published works (Ejaz, Hamada, and Diedrichsen 2015; Guan et al. 2022).

To intuitively visualize these features, the neural activity was projected into a three-dimensional space identified using five-dimensional cross-validated targeted dimensionality reduction (TDR) followed by PCA (see Methods). For each finger, neural activity corresponding to flexion and extension movements evolved along diametrically opposite directions from the origin, and nearby fingers occupied similar directions. In the top two PCs, thumb flexion overlapped with little finger extension, and thumb extension overlapped with little finger flexion (Figure 2F), explaining the negative correlation between thumb and little finger directions in Figure 2E. However, the neural activity for the thumb and little fingers were separated in the third PC dimension (Figure 2G), maintaining the ability to distinguish between them.

This structure was preserved across left and right hands and was not influenced by different palm orientations, with the limb-specific and/or pose-specific component linearly translating the activity in the neural state space (Supplementary figure 3, (Vargas-Irwin et al. 2022)).

### Neural activity for multi-finger movement is consistent with the sum of its parts

Given the neural representation of single finger movements, how are they combined to represent multi-finger movements? First, we confirmed that attempted multi-finger movements are robustly represented in our neural recordings. For a collection of 38 natural hand movements consisting of gestures from the American Sign Language (ASL) alphabet and some stereotypical single/multiple finger movements (Figure 3A), a linear decoder achieved 76% classification accuracy (chance 2.6%) using the neural activity recorded from T5 (Figure 3B). Since most of these gestures have correlated movements for nearby fingers and do not span the complete space of finger movements, we repeated this analysis with a collection of 80 “non-natural” combinatorial finger movements. For these combinatorial movements, each finger was independently cued to be either flexed, extended or at rest. To ensure that enough repetitions of each combination could be collected within a single research session, the ring and little fingers were linked together (referred to throughout as a single “finger” entity, the “ring-little finger”) to reduce the number of movement conditions to 80. A linear decoder achieved 40% classification accuracy (chance: 1.3%, Figure 3C). This shows that various natural and non-natural multi-finger movements are distinctly represented in the hand area of the premotor cortex, permitting a study of their neural geometry.

**Figure 3.**
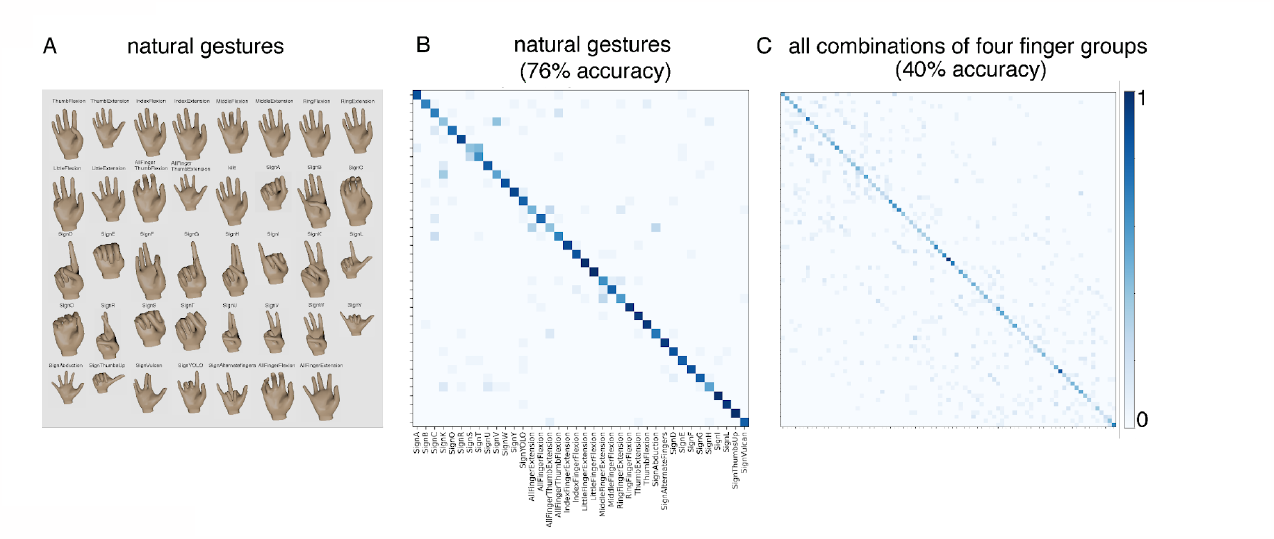
Attempted multi-finger movements are well-represented in the neural activity. (A) Neural activity was recorded as T5 attempted 38 hand movements on the right hand, consisting of gestures from the American Sign Language (ASL) alphabet & single finger movements. Each movement was attempted 22 times in trials consisting of 1 second prep, 1 second move and 2 second hold periods. (B) A confusion matrix, where the (i, j)th entry is colored by the percentage of trials where movement j was decoded when movement i was cued. A linear support vector classifier was used for decoding. Classification accuracy was 76%, substantially above the chance accuracy of 2.6%. (C) The same decoding analysis was applied to a collection of 80 combinatorial gestures, where four finger groups were varied independently (the ring and small fingers constrained to the same movement). Each finger group was independently cued to be flexed, extended or idle (15 trials were collected per condition, with a 1 second preparatory and a 2 second movement period). The classification accuracy was 40%, substantially above the chance accuracy of 1.3%.

Next, we tested the compositional coding hypothesis for multi-finger movements, beginning with the simultaneous movement of pairs of fingers. The neural subspace that was most modulated by each pairwise finger group movement was visualized with TDR after trial averaging and removing the condition invariant signal (Figure 4A, B, C). Activity for pairwise finger movements evolved in a region of the neural state space that was ‘in-between’ that of the corresponding single finger movements, as expected from linear summation of the constituent movements. Next, we assessed the degree of linearity quantitatively by fitting a linear summation model to a large set of 80 combinatorial finger movements, with four independently varying finger groups. The linear model learns a neural activity pattern for the flexion or extension of each finger, and then adds them together to predict the neural activity for a multi-finger movement. For each finger, we found that the magnitude of the representative activities was higher for flexion compared to extension; and for both types of movements, the magnitude was higher for thumb and ring-little fingers compared to index and middle fingers (Figure 4D). The linear model predicted neural activity significantly better than chance (Figure 4E) and captured 48% of the variance (Figure 5F).

**Figure 4:**
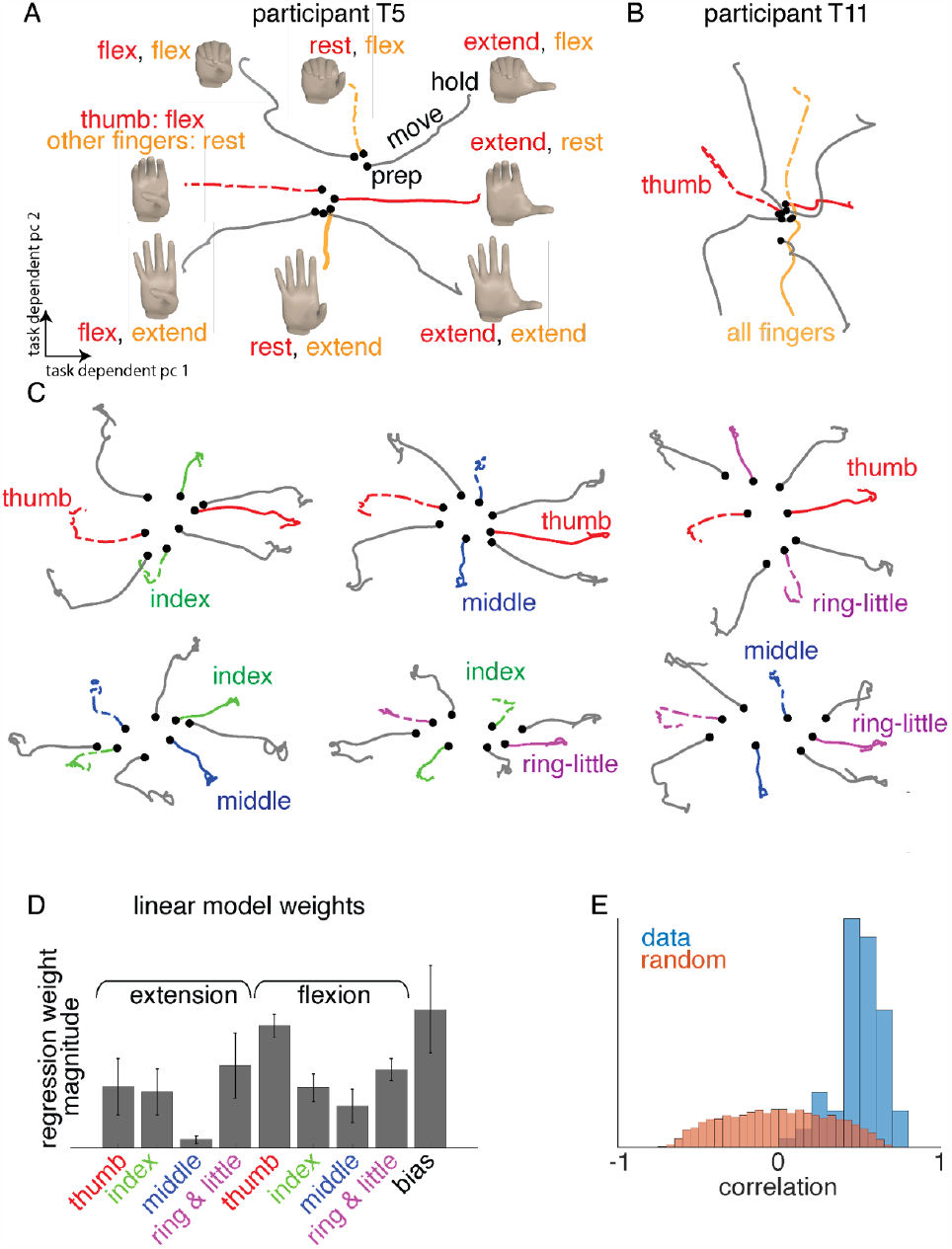
Neural activity for simultaneous movements evolves in a direction consistent with linear summation. (A) Participant T5’s neural activity for all movement combinations of two “finger groups” 1.the thumb, and 2. all other fingers linked together. For each movement condition, threshold crossings were trial averaged and concatenated (over the electrode dimension) across two sessions. Neural activity visualized after subtracting the condition invariant signal (CIS) and projecting-out the associated neural dimensions, followed by cross-validated targeted dimensionality reduction (TDR, see Methods). Lines indicate trajectories for each condition which start at rest, move outwards, and settle at a final position representing the target gesture. Trajectories where only one finger group moves are colored, while gray lines correspond to conditions where both groups move in combination (as shown by inset images). (B) Same as (A), using data from one session in participant T11 (see Methods). (C) Same as (A), using data from five sessions for pairwise combinations of four finger groups (thumb, index, middle and ring-little combined). (D) A linear encoding model was fit to predict trial-averaged neural activity recorded from T5 during the hold period (after removing the CIS) using the finger movement kinematics. Kinematics were represented with an eight dimensional one-hot encoding, with each dimension corresponding to the presence (=1) or the absence of (=0) of a combination of finger and movement type (flexion/extension). Bar heights indicate the magnitude of regression coefficients corresponding to each finger. Neural activity modulated more with flexion than extension, and thumb, ring & little, index and middle modulated activity in a decreasing order. (E) Distribution of the cross-validated correlation between observed neural activity and predictions from a linear encoding of kinematics (blue) with four dimensions (one dimension per finger, +1 for flexion, -1 for extension and 0 for rest). The linear model predicts the neural geometry better than the null distribution (orange) generated by randomly shuffling the neural activity and the kinematics labels with respect to each other and measuring the correlation with the linear model predictions.

**Figure 5.**
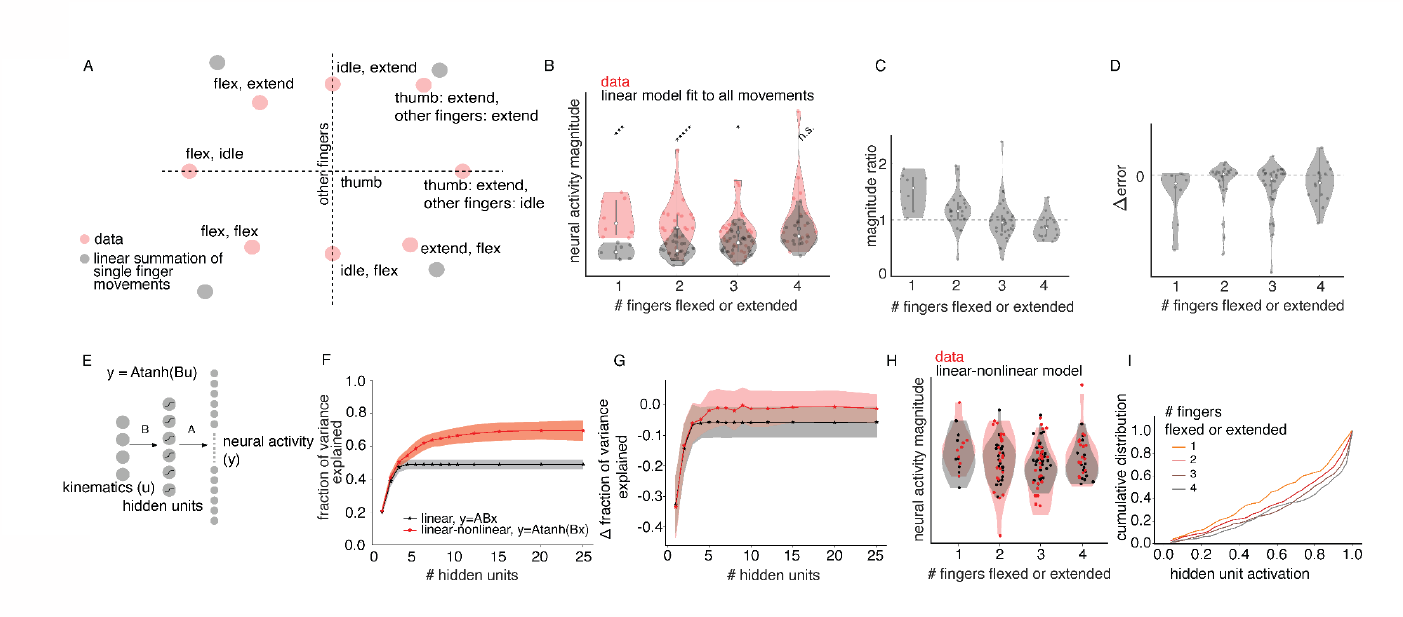
Overall neural activity magnitude is conserved for multi-finger movements. (A) Root mean squared magnitude of the recorded neural activity during the hold period after removing the condition invariant signal (red dots) for an example set of two finger-group movements (thumb vs. all other fingers). Distance from the origin corresponds to magnitude of neural activity, with different movement conditions represented along different directions. Predictions from linearly summing the single finger-group movements are indicated in gray. The magnitude of the recorded neural activity for two finger-group movements was lower than what would be expected from the sum of the corresponding single finger-group parts. (B) A comparison of recorded activity magnitude (red) and predictions from a linear model (black) for 80 combinatorial movements of four finger-groups. Movements grouped by the number of fingers either flexed or extended (x-axis). Linear model predicts that magnitude will increase as more fingers move, but the recorded activity magnitudes appear relatively constant. The linear model under-predicts the neural activity magnitude for a small number of moving finger-groups, as the linear model is fit to all movement conditions and there are a large number of conditions with multiple finger-groups moving. Statistical significance of difference in means evaluated with a t-test.(C) Ratio of the recorded activity magnitude vs. the linear model’s prediction. The ratio is clearly greater than 1 for single finger-group movements, showing that the linear model under fits the magnitude of neural activity. (D) Change in prediction error when a condition-dependent gain is added to the linear model – as expected, prediction error decreases for single-group movements. (E) A linear-nonlinear model which applies a tanh(.) non-linearity to map kinematics to predicted neural activity patterns; this nonlinearity should allow the model to capture activity magnitudes more accurately via saturation. *A* and *B* denote linear maps from kinematics to hidden unit inputs and hidden unit activity to the observed neural activity respectively. (F) Fraction of variance explained across the 80 combinatorial movements by the linear-nonlinear model with an increasing number of hidden layers (red). A low-rank linear model was used for comparison (gray). Linear-nonlinear models outperform the linear model and performance saturates at a small number of hidden dimensions of ∼10. Cross-validated performance was averaged across 100 resamplings of trials used for training and evaluation. (G) Similar analysis as (F), but testing generalization across movements – models were trained on a random subset of movements and tested on the remaining movements. Linear-nonlinear model generalizes better than a linear model to novel movement conditions. Since the performance varies based on which movements are partitioned into the test set, the performance change was measured compared to the best model within each partitioning (linear-non linear model with the highest number of hidden units), and changes were averaged across resamplings. (H) The linear-nonlinear model (with 20 hidden units) better captures the magnitude of neural activity (compared to B). Difference in means was not statistically significant (t-test). (I) Cumulative distribution of hidden unit activity in the linear-nonlinear model for conditions with different numbers of moving finger-groups. Note the greater saturation with a larger number of moving finger-groups.

### Overall neural activity magnitude is conserved for multi-finger movements

While the linear summation model is simple and intuitive, it makes a rather strong prediction: that the magnitude of neural activity should increase with the number of moving fingers (i.e., the total number of fingers either flexed or extended). We tested this prediction by comparing the magnitude of the neural activity to that predicted by a linear model. We measured the magnitude of the neural activity during the hold period (avoiding potential trial-to-trial variability in reaction time and movement speed), using an unbiased estimate of activity magnitude (Willett et al.2020). For two finger-group movements, the magnitude of neural activity for simultaneous movement was smaller than what would be expected from summing the constituent single finger-group movements (Figure 5A shows an example of two finger-groups, Supplementary Figure 4 shows others). For combinatorial movements of four finger-groups, neural activity magnitude remained constant as the number of moving fingers increased. A linear model fit to all movement conditions failed to capture this phenomenon, and instead predicted that magnitude should increase with the number of moving fingers. This resulted in an under-prediction of neural activity magnitude for single and two finger group movements (Figure 5B). In sum, the observed magnitudes are consistent with normalization, and a linear summation model fails to capture this.

A simple extension of the linear model, where the predictions are simply multiplied by a movement-specific scalar, showed an improvement in the variance explained for single-finger movements (Figure 5C-D), with the scale being greater than one (Figure 5C). While the scaled linear model captures the observed normalization effects, it requires a condition-specific scalar and does not generalize from one subset of movements to another subset of movements.

We tested a flexible model without movement-specific parameters, which can achieve normalization of neural activity through saturating nonlinearities in the mapping from kinematics to neural activity. Neural activity (*y*, during the hold period) is modeled in terms of kinematics (*u*) using a *pseudo-linear model*: *y* = *A tanh* (*Bu*), where *B* defines a linear map from kinematics to the hidden units and *A* defines a linear map from the hidden units to the neural activity (Figure 5E). The kinematics vector *u* encodes the positions of the four finger groups in a binary fashion, with +1/0/-1 corresponding to flexion/rest/extension. This model can be interpreted as explaining the neural activity by linearly combining the kinematic variables and then non-linearly projecting it onto a lower-dimensional manifold described by the hidden unit activities. The linear-nonlinear model outperformed the linear model when trained on an identical set of movement conditions as test data (Figure 5F) and could generalize successfully across movements (test movements different from train movements, Figure 5G). Additionally, the performance of the linear-nonlinear model is saturated with ∼10 hidden units, much smaller than the maximum number of hidden units that may be needed (= minimum of number of movement conditions (80) and the dimensionality of neural state space (448)). This implies that the neural activity patterns themselves are low-dimensional, and that therefore multi-finger movements largely exist along directions of the neural space already explored by single-finger movements. Finally, the linear-nonlinear model captured magnitude normalization, i.e., how neural activity remained at a relatively constant magnitude with an increasing number of moving fingers (Figure 5H). The model achieved this via saturating nonlinearity, as shown by the greater saturation of hidden unit activity with increasing number of moving finger groups (Figure 5I).

### Pseudo-linear representation limits linear decodability of fingers with weaker modulation

What is the impact of pseudo-linear compositional representation on the ability to use a linear decoder to identify the finger positions during multi-finger movements? This can be assessed by measuring how the representation of a given finger changes across contexts, i.e., how it changes with particular movement combinations of the other fingers. In the neural state space, the line connecting a pair of multi-finger movements which only differ in the final position of a target finger (i.e., either flexed or extended) but which are identical in the final position of other fingers describes the target finger’s ‘population tuning’, i.e., how the population activity changes as a function of a finger’s movement in a given context. Changes in the population tuning across contexts measures how a decoder might generalize to multi-finger movement it was not trained on. A linear decoder, for example, would decode the motion of each finger by projecting neural activity along each finger’s population tuning direction – in this case, generalization would be impaired if the population tuning changes.

Three possible outcomes could describe how the neural population tuning could change as a function of context (Figure 6A, B, C). If the contribution of individual fingers is largely summed linearly to represent multiple finger movements, then there would be no change in the population tuning magnitude and direction across contexts (Figure 6A). Hence, a linear decoder that can identify the finger position in one context could generalize to other contexts. However, if multi-finger movements result in changes in tuning magnitude and/or tuning direction, a linear decoder for a given finger would not generalize optimally across contexts (Figure 6B, C).

**Figure 6.**
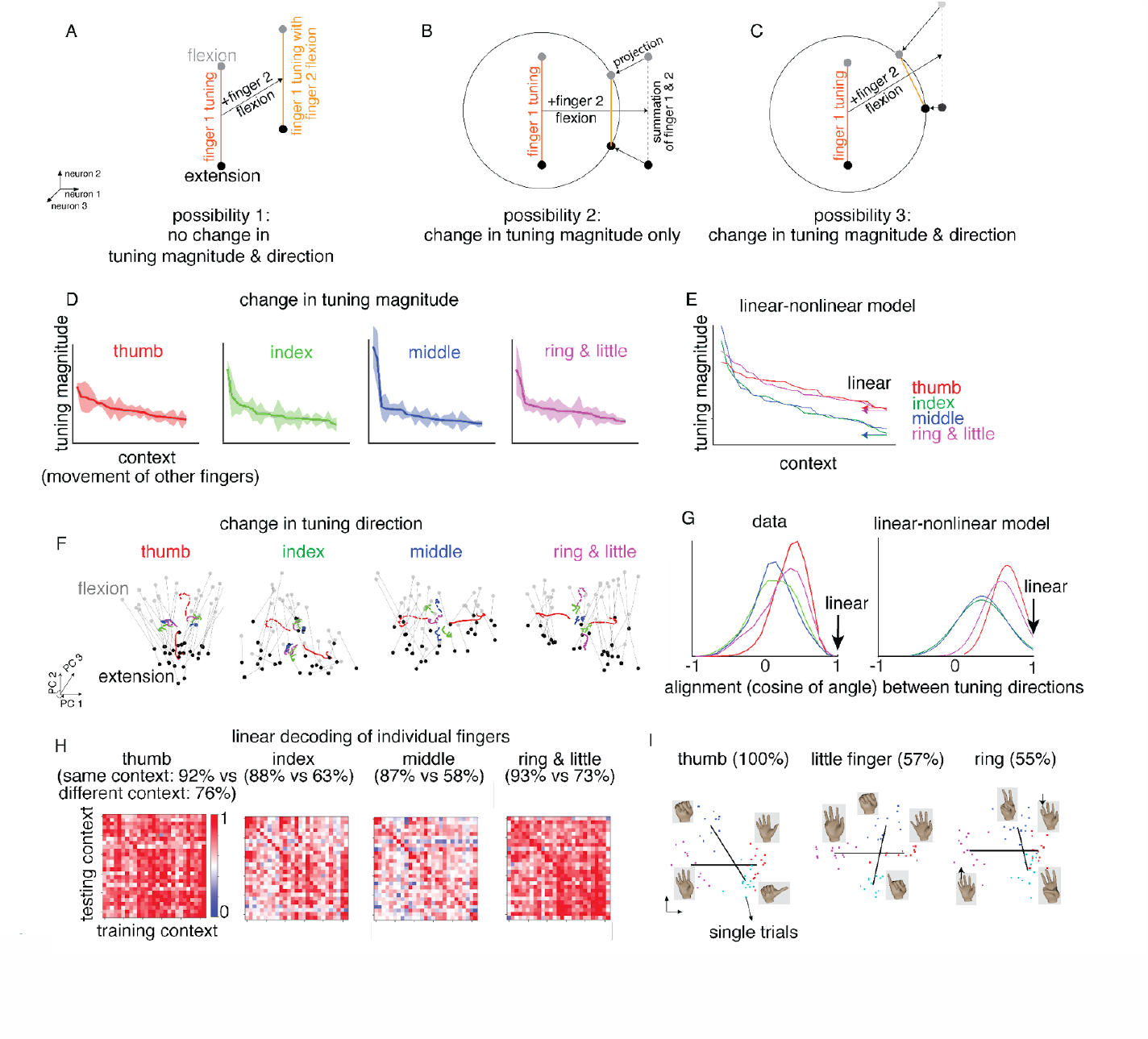
Impact of pseudo-linear representation on the linear decodability for individual fingers. (A,B,C) Different ways in which the linear decodability of a finger (finger 1) could be affected by the simultaneous movement of another finger (finger 2). Pseudo-linear combination is indicated in the neural state space by linear summation of neural activities and projection of the resulting activity onto a unit-circle. An optimal linear decoder for the position of a finger projects neural activity along the line joining the points corresponding to the flexion and extension movements (‘population tuning’). Comparing the population tuning for finger 1 during the isolated movement and simultaneous movement with finger 2, we could either expect no change in direction and magnitude (A, would be expected from pure linear summation), change in magnitude only (B, would be expected from normalization combined with orthogonal tuning directions of finger 1 and 2), or change in both direction and magnitude (C, caused by normalization combined with correlated tuning directions). (D) How neural population tuning magnitude for a given finger group is affected by the movement of other fingers. Data are shown from 80 combinatorial movements of four finger groups. “Contexts” (specific movements of other fingers) are sorted by tuning magnitude. (E) Context dependence of population tuning magnitude in a linear-nonlinear model from Figure 5E. While the linear-nonlinear model captures the variations in tuning magnitude, a linear model has constant tuning magnitude by construction (arrows). (F) How is the population tuning direction for a finger affected by the movement of other fingers? Three principal components capturing the average neural activity for flexion-extension movements across fingers are computed and visualized with a two dimensional projection that aligns the flexion-extension movements of a particular finger along the y-axis. Colored lines indicate the marginalized (average across conditions) neural activity for flexion (dotted) and extension (solid) movements of fingers. Gray (black) dots indicate individual conditions where the target finger was flexed (extended). Lines join movement pairs from the same context (i.e., other fingers have the same movement). Tuning directions (line angles) change depending on the context. (G) Histogram of alignment of population tuning directions (cosine of angle) across pairs of contexts in data (left) and linear-nonlinear model fits (right). Population tuning directions are more aligned for thumb and ring/little finger groups in both data and the linear-nonlinear model. By construction, the linear model exhibits complete alignment of all contexts (all values at 1). (H) Performance of a linear decoder (support vector classifier) when classifying between flexion/extension positions of a finger across contexts. Element at position (i,j) corresponds to training the classifier on context j and testing on context i. Mean within-context (diagonal values) accuracy was high for all finger groups, whereas across-context (off-diagonal values) accuracy was low for middle and index fingers compared to the thumb and ring-little group. (I) Examples of cross-context decoding performance for the thumb, little and ring finger. Two gesture pairs are shown for each finger. For the first pair, the target finger is either flexed or extended while the other fingers are at rest. For the second pair, the target finger is either flexed or extended while all other fingers have an identical movement. A linear classifier for thumb position trained on isolated movements showed 100% accuracy when tested on the second pair of movements (Thumbs Up vs ASL sign for letter S). This accuracy dropped to nearly chance performance in a similar analysis for the little and ring fingers.

We evaluated these possibilities by measuring the population tuning magnitude and direction for each finger across contexts using neural activity for combinatorial movements of four fingers. For each finger, there was a context-dependent change in tuning magnitude (Figure 6D), with a greater variation for the index and middle fingers compared to the thumb and the ring-little finger. Similarly, tuning directions varied more for the index and middle fingers (Figure 6F, G, Supplementary movie 1). While a linear model, by definition, cannot describe changes in tuning direction or magnitude across contexts, the linear-nonlinear model proposed above captured both of these properties (Figure 6E, G). Overall, this suggests that fingers which are weakly represented in the neural activity (ex. index and middle fingers) show a greater change in population tuning during multi-finger movement. Intuitively, this can be attributed to the saturating effect of magnitude normalization which results in a loss of weakly represented information but preservation of strongly represented information.

We assessed the impact of these population tuning changes on the generalizability of linear decoding across contexts using two tasks. For combinatorial movements of four finger groups, the within-context decoding performance was high across all fingers (88-92%); across-context performance was much lower for middle and index fingers (58% and 63% respectively) compared to thumb and ring-little fingers (73% and 76% respectively) (Figure 6H).

These observations were replicated for pairs of natural gestures. Specifically, a linear decoder trained on isolated flexion/extension movements of the thumb generalized to gesture pairs where all other fingers were flexed but the thumb was either extended (‘thumbs up’ gesture) or flexed (American Sign Language for symbol S, Figure 6I). However, for other gesture pairs, a linear decoder trained on isolated movements of the ring finger or the little finger performed poorly (Figure 6I), suggesting that the population tuning for the movement of ring and little finger separately changes more compared to moving them as a finger group. As a consequence of these context-dependent changes in population tuning, non-linear decoders such as a recurrent neural network and a temporal convolutional network performed better than linear (Kalman) filters for decoding fingers positions, with a larger difference in performance for index and middle fingers (see Supplementary Figure 5).

In summary, the population tuning of weakly-represented fingers changed more based on the movement of other fingers, suggesting that peak performance for iBCI finger control requires the use of non-linear decoders that jointly decode the movement of all fingers.

## Discussion

We discovered a pseudo-linear compositional neural code for how individual finger movements are combined to represent multi-finger movements. While the neural activity evolved in a direction consistent with the linear summation of its constituent movements (Figure 4), the magnitude of neural activity was normalized, i.e., it was independent of the number of moving fingers (Figure 5). As a consequence, fingers that were weakly represented (such as middle) showed a greater change in their tuning magnitude and direction as a function of the movement of other fingers (Figure 6). Hence, while the linear model explained a large fraction of the variance in the neural activity, decoding multiple finger movements can be substantially improved with a joint, non-linear decoder (Supplementary Figure 5).

The pseudo-linear composition of individual finger movements is consistent with recent work in monkeys (Nason et al. 2021), which showed that linear summation of neural activity for single finger movement explains two-finger movements. However, our analysis of the two finger and four finger-group movements (enabled by the ability of human participants to perform complex and flexible tasks) revealed a characteristic deviation from a pure linear summation. While the dominant fingers (e.g., thumb) had consistent tuning across different combinations (as predicted with linear summation), the weakly-represented fingers (e.g., middle) showed a larger deviation from the linear model. This is explained with magnitude normalization after linear summation, and is consistent with two recent publications. First, prior work in monkeys showed that in a task requiring production of different amounts of force, a linear model needs to be supplemented by a condition-dependent gain to predict EMG activity from neural activity (Naufel et al. 2019). Second, there exists a mild degree of non-linear encoding for two finger movements in monkeys, explaining why non-linear decoders have been shown to improve closed loop performance (Willsey et al. 2021).

In addition to finger movements, pseudo-linear composition may explain the neural activity for simultaneous limb and multi-joint movements as well (Cisek, Crammond, and Kalaska 2003; Rokni et al. 2003; Diedrichsen, Wiestler, and Krakauer 2013; Wiestler, Waters-Metenier, and Diedrichsen 2014; Yile Jin et al. 2016; Fujiwara et al. 2017; Bundy et al. 2018; Ames and Churchland 2019; Willett et al. 2020; Cross et al. 2020; Downey et al. 2020; Dixon et al. 2021; Merrick et al. 2022). Future work will verify if magnitude normalization can explain the observed suppression of the non-dominant limb for simultaneous bimanual movements and if there exists a unified theory of how the brain represents simultaneous multiple movements.

What circuit mechanisms could explain the mild degree of nonlinearity and magnitude normalization? One hypothesis is that the observed non-linearities could be a consequence of inputs with different strengths processed by a network of neurons with saturating nonlinearities. We tested if a simple recurrent neural network (RNN) with static input drive of unequal magnitudes and saturating unit activations could capture the observed properties. The inputs correspond to different fingers, and the unequal input magnitudes reflect the unequal modulation observed for different fingers. Starting from random unit activation, the RNN was simulated until the activity reached a steady state. This steady state activity captured all the properties observed for neural recordings – it was low dimensional, with a largely constant magnitude that cannot be approximated by a linear model of finger positions, and the population tuning direction for weaker inputs changed more compared to the stronger inputs (Supplementary Figure 6).

This work helps to address a fundamental question of how neuronal ensembles represent multiple task-related variables. While a neural representation that linearly composes individual task variables is simple, and a decoder trained to decode a particular task variable in one context generalizes to other contexts, the overall capacity to accurately represent multiple combinations of task variables is limited. An alternative representation using non-linear composition of task variables provides a higher capacity but poorer decoder generalization across contexts. Similar to the pseudo-linear compositional principle described here, several studies have presented evidence of a largely linear composition with a mild non-linearity - a compromise between the two extremes. (Saxena and Cunningham 2019; Bernardi et al. 2020; Chung and Abbott 2021; Nogueira et al. 2023). However, most of the literature on multi-task computation has been limited to simulated networks (Yang et al. 2019; Driscoll, Shenoy, and Sussillo 2022), neural recordings in non-human primates (Bernardi et al. 2020; Xie et al. 2022), or functional magnetic resonance imaging in humans (Ito and Murray 2022). Along with some recent works (Willett et al. 2020; Guan et al. 2023), we extend the principle of compositionality to neural recordings in the human motor cortex at single neuron resolution, providing further evidence that compositionality may be a general feature of brain representations.

Overall, we show that the compositional coding hypothesis with pseudo linear summation (Figure 1B) is strongly supported by the neural geometry of multi-finger movements in the human premotor cortex. This principle may explain the neural structure in other motor and cortical tasks and may aid in a rapid development of future BCI applications such as multi-finger typing, playing a piano or whole-body movement.

## Acknowledgement

The authors would like to thank T5, T11 and their care partners. We would like to thank David Sussillo for helpful discussions. We would like to thank Beverly Davis, Kathy Tsou, and Sandrin Kosasih for administrative support.

Support provided by Office of Research and Development, Rehabilitation R&D Service, Department of Veterans Affairs (N2864C, A2295R); Wu Tsai Neurosciences Institute; Howard Hughes Medical Institute; Larry and Pamela Garlick; Samuel and Betsy Reeves; Sons Foundation Collaboration on the Global Brain 543045; NIDCD R01-DC014034, NIDCD U01-DC017844, NINDS UH2-NS095548, NINDS

U01-NS098968.

* The contents do not represent the views of the Department of Veterans Affairs or the US Government. CAUTION: Investigational Device. Limited by Federal Law to Investigational Use

## Methods

### Participant information

Research sessions were conducted with volunteer participants enrolled in the BrainGate2 pilot clinical trial (ClinicalTrials.gov Identifier: NCT00912041). The trial is approved by the U.S. Food and Drug Administration under an Investigational Device Exemption (#G090003, Caution: Investigational device. Limited by Federal law to investigational use) and the Institutional Review Boards of Stanford University Medical Center (protocol #20804), Brown University (#0809992560), and Partners HealthCare / Massachusetts General Hospital ((#2009P000505).

Participant T5 is a right-handed man who was 69 years old at the time of the study. He was diagnosed with C4 AIS-C spinal cord injury eleven years prior to this study. T5 is able to speak and move his head, and has residual movement of his left bicep as well as trace movement in most muscle groups (See (Willett et al. 2020) for further details). T5 gave informed consent for this research and associated publications.

Participant T11 is a 36-year-old right-handed man with a history of tetraplegia secondary to C4 AIS-Aspinal cord injury that occurred 11 years prior. T11 gave informed consent for this research and associated publications.

Both participants had two 96-channel intracortical microelectrode arrays placed chronically into the hand knob area of the left precentral gyrus (PCG).

### Neural recordings

For the present study, neural control and task cuing closely followed (Gilja et al. 2015) and were controlled by custom software running on the Simulink/xPC real-time platform (The Mathworks, Natick, MA), enabling millisecond-timing precision for all computations. Neural data were collected by the NeuroPort System (Blackrock Microsystems, Salt Lake City, UT) and available to the real-time system with 5 ms latency. Neural signals were analog filtered from 0.3 Hz to 7.5 kHz and digitized at 30 kHz (250 nV resolution). Next, a common average reference filter was applied that subtracted the average signal across the array from every electrode in order to reduce common mode noise. Finally, a digital bandpass filter from 250 to 3000 Hz was applied to each electrode before spike detection. For threshold crossing detection, we used a −4.5 x RMS threshold applied to each electrode over 1ms bins, where RMS is the electrode-specific root mean square (standard deviation) of the voltage time series recorded on that electrode and downsampled at 15 or 20 ms bins. For most analyses the binned activity was also smoothened with a gaussian filter (σ = 300ms).

For T11 data, neural activity was recorded using the Brown Wireless Device (BWD) described previously (Simeral et al. 2021). BWD transmitted data at 20 kHz and 12 bits per sample, which was then upsampled to 30kHz and 16 bits per sample. The raw activity was then passed through a fourth order bandpass butterworth filter (250Hz-7500 Hz). A common average reference filter was applied, threshold crossing counts were computed for 1 ms bins (threshold = -4 x RMS) and discretized at 15 or 20 ms bins and Z-scored (normalized) for each channel.

### Task implementation

The finger and hand visualization was developed in Unity Software (Unity Technologies, San Francisco) using a pre-fabricated hand (Barnett, n.d.). The fingers and hand were animated by either using the Animation toolbox in Unity to continuously interpolate between specified starting and ending positions (American Sign Language gestures), or specifying the trajectory of joint positions and rotations directly from an external program using Redis (Redis Enterprise, all other tasks, which have a single axis of motion per finger). When specified directly, motion involved only the flexion-extension movements of individual fingers and joint positions and angles were interpolated between a complete flexion and extension position.

All blocks were ‘open-loop’ and consisted of an animated hand, smoothly transitioning from one gesture to another, and the participant was asked to attempt to make those same motions with his own hand even though paralysis prevented him from doing so. Each trial consisted of distinct phases: a preparatory period, a go cue followed by a movement period, and a hold period (optional). During the preparatory period, participants were shown the target finger positions but did not move their hands. At the go cue, they were instructed to slowly move their hands to the target positions. During the hold period, they were instructed to hold their hands steady at the target positions.

The trial structure followed a structure similar to the 2D cursor control task common in motor neuroscience (Georgopoulos, Kettner, and Schwartz 1988)). Borrowing the terminology from 2D cursor control tasks, trials alternated between ‘center-out’ and ‘return’ trials. Center-out trials began with the hand at a rest position, which then proceeded to move to a target gesture (equivalent to the “going-out” phase of a center-out cursor control task) . During return trials, the hand moved back to the rest posture. Only center-out trials were analyzed.

For the single finger movement task, a single finger was either fully flexed or extended in each trial. Each trial consisted of a 1.5s preparatory phase and a 1s movement phase. Each block consisted of 81 gestures (162 trial total as center-out and return trials alternate). Data from this task was analyzed in Supplementary Figure 3.

Center-out single finger movements were performed across contexts (different hands and palm-poses), either changing contexts across blocks (but fixing the context within each block) or alternating across contexts within a block. Data from this task was analyzed in Figure 2.

For combinatorial movements of four finger-groups (the ring and little fingers were constrained to move together), the target position for each finger-group was sampled independently (full flexion/full extension/rest), resulting in 81 unique movements, with one movement being rest. These 81 movements were randomly divided into three groups consisting of 27 movements each. Each block only contained movements from one of these groups, and each movement was repeated three times in a row. The multiple repetitions of each movement were designed to improve behavioral reproducibility, since the movements were often non-intuitive and non-natural. Hence three open-loop blocks with different subsets of target configurations give three trials of all 81 target gestures. Each trial had 1s preparatory time and 2s move time (slightly longer move time reduces behavioral variability). Data from this task was analyzed in Figure 3,4,5,6.

Sign-language task consisted of 38 total gestures, with some corresponding to signs from the American Sign language and others added probe single-finger movements, coordinated movements of all fingers (e.g. all finger flexion/extension) or decorrelating the movements of all fingers (e.g. Sign Alternating Fingers). Each trial had 1s preparatory time, 1.5s move time and 2s hold time. Data from this task was analyzed in Figure 3.

### Neural geometry analysis

#### Unbiased estimate of neural magnitude

The unbiased estimator outlined in (Willett et al. 2020) was used to estimate neural magnitudes and distances. First, we outline why a “naive” estimator based on sample estimates (instead of cross-validation) is biased. For trial *i* of a given movement, let *n*_*i*_ be the vector of recorded neural activity across channels with mean *n*. Let the noise be additive, zero-mean and independent across trials (*ϵ*_*i*_), giving *n* _*i*_ = *n* + *ϵ*_*i*_. A naive approach to estimate the magnitude (||*n*||^2^) would be to estimate the magnitude of the sample mean (trial average): 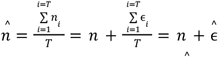. The magnitude of this estimate is 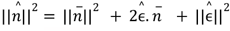. Since the expected value of 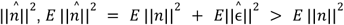 is zero, the second term is zero. However, 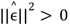 making the expected value of 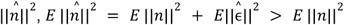 showing that the naive estimate is biased.

An unbiased estimator of the magnitude ξ(||*n*||), *n*^2^) can be achieved as follows. Let us define 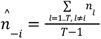 the empirical mean of all trials except the *i*th trial. An unbiased estimator can then be defined using disjoint dot products as follows: 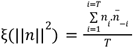. Note that, 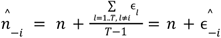. The expected value of each term in the summation is given by:

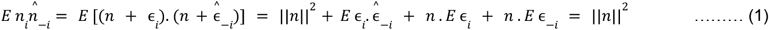

Since the noise is zero mean and independent across trials, the last three terms evaluate to zero, making the summation unbiased.

When using data from multiple sessions, the unbiased estimator is applied on all trials within each session first; the session-specific estimates are then averaged across sessions. Note that the unbiased estimator could give negative values, which we interpret as evidence that the true magnitude is near zero.

#### Representational dissimilarity matrix analysis using an unbiased estimate of neural distances (Figure 1)

A representational dissimilarity matrix is constructed by calculating the neural distance between all pairs of movement conditions. A naive estimator that computes Euclidean distances directly between trial-averaged activity vectors is biased, as explained above. An unbiased estimator of Euclidean distance can be achieved by using the unbiased estimator of neural magnitudes for individual conditions. Let *n*^*a*^ and *n*^*b*^ correspond to the mean activity vectors for distributions of activity observed for conditions *a* and *b*. Using the mathematical identity ||*n*^*a*^ − *n*^*b*^ ||^2^=||*n*^*a*^||^2^+ ||*n*^*b*^||^2^− 2*n*^*a*^. *n*^*b*^, the neural distance estimator is given by

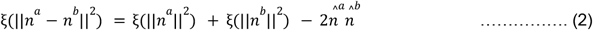

exploiting the unbiasedness of the first two terms (from above) and the independence of noise across trials. If *n*^*b*^ is deterministic (not random), the second term simplifies to || *n*^*b*^|| ^2^.

#### Unbiased estimate of neural magnitudes under linear summation (Figure 5)

Similar to the estimate of distances, the magnitude of neural activity when two neural activity patterns are added together (||*n*^*a*^ + *n* ^*b*^||^2^) is given by

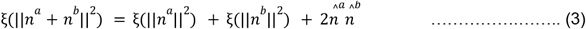

#### Marginalized principal component analysis (Figure 6F)

The visualization of neural activity in Figure 6F aims to isolate the contribution of each finger group during the simultaneous movement of the four finger groups (thumb, index, middle and ring-little constrained together). The motivation is to visualize the schematic from Figure 1A, B, C using real data. Briefly, we identify a three dimensional subspace that captures the activity across all single finger movements, and visualize the activity in this subspace after aligning the tuning directions for a particular finger to be vertical.

Each finger was independently flexed, extended or was constrained to be idle, resulting in 80 combinations (3^4^ = 81, with the movement with all fingers at rest removed). Neural activity (C=192 channels) was recorded in T5 across five sessions. Neural activity for each trial was binned and smoothened (details above) resulting in T=130 samples per trial. The analysis in this section used the trial-averaged activity for each movement condition. The trial-averaged activity was concatenated across sessions, resulting in C (=192, number of channels) x S (=5, number of sessions) dimensional observations. Trial-averaging within a session minimizes noise correlation between channels and concatenating the channels across sessions is justified if we assume that we get a random readout of the same underlying task-related manifold for each session. Let the trial-averaged and session-concatenated activity for movement condition *i* be represented by *T* ×*CS* dimensional matrix *n*_*i*_.

First, condition invariant signal is calculated by averaging the activity across all movement conditions 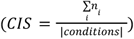. A rank-3 approximation of CIS is computed using singular value decomposition, and all *n*_*i*_ are projected orthogonal to it.

Next, the contribution of each finger movement is computed by marginalizing-out the movements of other fingers. For example, the contribution of thumb-flexion is calculated as

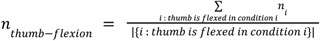 and is of dimensions *T* ×*CS*. Next, these marginalizations are concatenated for the flexion/extension movements on each finger and all four finger groups and to give 8*T* ×*CS* dimensional matrix. Projection onto a top three PCs (reducing the matrix to 8*T* ×3) gives the 3D visualization of all finger-movement combinations. Further, this space can be rotated to align flexion-extension movement of each finger in the north-south direction. Dots are generated by projecting the neural activity of each movement combination (*n*_*i*_) onto this 3D space and calculating the average of the last quarter of the trial duration (presumably corresponding to holding a given gesture). Only conditions when a particular finger is not idle are considered, with black dots corresponding to extension and gray for flexion; lines join condition pairs where other fingers have the same movement.

#### Cross-validated targeted dimensionality reduction

Targeted dimensionality reduction (Mante et al. 2013) projects the neural activity into a subspace that can be linearly decoded from movement kinematics. TDR projections from different partitions are aligned before averaging the projections of test data. The procedure is detailed below.

Let *n*_*s*_ be *N*_*s*_ ×*T*×*C* dimensional the neural activity for research session *s* with *N*_*s*_ trials, trial length *T* and *C* channels. Each neural activity pattern is associated with a kinematic movement condition *m*_*s*_ of *K* dimensions. The following steps are repeated multiple times:

1. *Partition data:* For each session and movement condition, the trials are partitioned into *training* and *testing* samples (random 50% split for each combination of session and movement condition).
2. *Marginalize:* The training data are trial averaged for each movement condition; time-averaged for the last section of the trial corresponding to continuously holding a given finger position; and concatenated across sessions. Hence 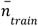 is *SC* ×*L* dimensional, with L being the total number of conditions. The test trials are similarly marginalized, but the temporal dimension is not collapsed, to give 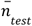 of dimension *SC* ×*T*×*L* .
3. *Remove condition-invariant signal:* Movement condition-invariant component is estimated from the training data and removed from 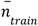 and 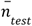. The movement condition-invariant neural activity is estimated by averaging all training neural activity (not averaged over time) and, concatenating it across sessions to give *n*_*cis*_ of dimensions *T*×*SC*. A low (three)-dimensional approximation *n*_*cis*_ is computed and 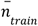 and 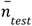 are projected orthogonal to this space.
4. *Linear regression:* The movement kinematics are encoded in a a (*K* + 1)×*L* matrix M, with each column corresponding to a different movement condition, and the first K rows correspond to positions of the fingers and the last row =1 is for estimating a bias. Finger positions are encoded as +1 for flexion, -1 for extension and 0 for rest. The number of non-zero elements in each column (excluding the last element corresponding to bias) indicates the number of fingers not at rest. The encoding matrix *A* of dimensions *SC* ×*K* is estimated by solving 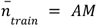 = *AM* using ordinary least squares. Top three left singular vectors from the SVD of *A* are retained for projecting neural activity.
5. *Alignment (for repetition > 1)*:The projection direction can vary across repetitions of this procedure. In the absence of any estimation noise, the projection directions can be sign-flipped across repetitions. In the presence of estimation noise, the projection directions can also be permuted or more generally, rotated across repetitions. While we do not attempt to correct for the rotations, we attempt to correct for permutations and sign-flips by aligning the projections estimated in later repetitions (> 1) to the first repetition by greedily matching the absolute value of the inner-product between projection directions.
6. *Projection:* The non-time averaged test data 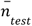 is projected onto the three dimensional space and averaged with previous repetitions.

This method was used in Figures 2F, G and Figures 4 A-C. The entire procedure is repeated 500 times for Figure 2F, G and 100 times for Figure 4A-C. Three sessions were used for Figure 2F, G five sessions for Figure 4A, C; and one session for Figure 4B.

### Encoding models for predicting neural activity from movement kinematics

Encoding models predict the neural activity observed for a given movement pattern. Neural activity corresponding to 80 conditions resulting from independent idle/flexion/extension movements of four finger groups (thumb, index, middle and ring-little) from four sessions were used. Neural activity was thresholded, binned, smoothened (details above), and only channels with high mean activity were kept (mean firing rate > 1.3Hz).

Trials were split into a disjoint training set (60% of the trials) and testing set (40%) for each session and condition. Condition invariant signal (average of the neural activity for different movements) was calculated for each session from the training data and subtracted from both training and testing data. Trial averaged and time-averaged (for the last quarter of trial duration corresponding to hold period) neural activity was estimated for each condition in both training and testing data. The magnitude for CIS-subtracted data was calculated using Equation (2) above. Movement kinematics were either represented as a one-hot vector with separate dimensions for flexion and extension, resulting in total number of dimensions equal to twice the number of finger groups (Figure 4D), or each finger group represented by either +1 (extension), 0 (rest) or -1 (flexion), resulting in total number of dimensions equal to the number of finger groups (other figures). In addition to splitting the trials, the cross-condition generalization analysis (Figure 5G) involved splitting the 80 movements into training (80%) and testing (20%) conditions.

#### Linear encoding model

A linear model approximated observed neural activity as a linear function of movement kinematics: *n*_*i*_ = *Ak*_*i*_ + *a*, where *n*_*i*_ is the neural activity vector for condition *i, k*_*i*_ is the kinematics, *A, a* are the learned encoding matrix and the bias respectively. Performance on testing data was calculated using MSE, estimated with Equation 2 (with one random and one deterministic component)) above.

For neural magnitude comparison in Figure 5B, the neural activity was 960 dimensional after concatenating trial averaged activity across five sessions and fit to the training partition consisting of 50% data. Condition Invariant Signal was estimated from training data and subtracted from both training and testing partitions. Unbiased estimate of neural activity was measured from test data. Conditions were grouped according to the number of moving fingers (either flexed or extended) and a t-test compared the distribution of data and model predictions for each group of conditions.

Optionally, a condition-dependent scalar *s*_*i*_ was estimated, that uniformly scales the all the dimensions of the neural activity to give encoding model: *n*_*i*_ = *s*_*i*_ (*Ak*_*i*_ + *b*). The estimated scales and corresponding performance improvement is shown in Figure 5C, D. Scales estimated from 95 resamplings of training (70%) and testing (30%) of data, across the four sessions, which had at least 8 trials per condition.

#### Linear non-linear encoding model

The linear model was augmented by saturating nonlinearities in a low dimensional space giving the encoding model: *n*_*i*_ = *B tanh*(*Ak*_*i*_ + *a*) + *b*. The parameters of this model (*A, a, B, b*) were optimized for minimizing an unbiased estimate of the MSE (Equation 2) using Adam optimizer (learning rate = 0.0001, stopped when change in loss < 1e-7).

In Figure 5F, the performance was compared for the linear-nonlinear model and a low-rank linear model with increasing hidden layers. Both models were trained on 70% of data and tested on the remaining data. All conditions used for training and testing. Performance was averaged over 95 resamples of data.

In Figure 5G, comparison of across-condition generalization performance for linear-nonlinear and linear encoding models. Training was performed on 80% of conditions (out of 80 total) and testing was performed on the rest of the conditions. Performance was averaged over 95 resamplings. Performance difference from the best model (linear non-linear model, with the largest number of hidden dimensions) indicated on y-axis.

##### Recurrent Neural Network (RNN) simulation

A mechanistic explanation for the linear-nonlinear encoding we observed was explored using simulations of a randomly connected recurrent neural network (Supplementary Figure 6). A recurrently connected network of *n* = 1000 neurons with tanh nonlinearity were simulated, with random connectivity *J* between them (connection between neurons *i* and *j, J*_*i,j*_ ∼ N(0, 1/n)). Two inputs (*i*_1_, *i*_2_) were applied to the network and varied between − 10 and 10. Each input was projected onto the neurons using input weight coefficients sampled from an identical distribution (vector *b*_*l*_ of length 1000), but the inputs had different scales (*s*_1_ = 5, *s*_2_ = 1).

For each input combination in a uniformly sampled grid of the two inputs, network activity was simulated from a random initial state until the activity settled to a steady state (magnitude of change in state < 1*e* − 6). The following update equation was used with τ = 1 and discretization step size 0.01.

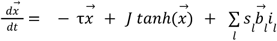

Analyses applied to the simulated data in Supplementary Figure 6 were the same as those we applied to the real data.

### Discrete decoding of movement kinematics from neural activity

Classification accuracy between a set of discrete movements is reported in Figure 3B, C and Figure 6H, I. To generate these classification accuracies, threshold crossings were first binned (10ms bins) and smoothed (by convolving with a gaussian kernel, sd=20ms). Binned threshold crossing rates were then averaged across time within a 1.5s window (entire duration of the go period), yielding a single neural feature vector for each trial. inactive channels (as defined by <10 Hz activity) were then removed from the feature vectors, activity for each channel was Z-scored within each block.

The ability to distinguish between different gestures was assessed using a linear classifier (support vector classifier from the sklearn package (Pedregosa et al. 2012), linear kernel, C=0.025). For the analysis in Figure 6H and Figure 6I, classifiers were trained on one pair of gestures and evaluated on different pairs. For 6I, instead of using the mean activity for the entire duration of the trial, the accuracy is reported with a featurization of neural activity corresponding to a concatenation over 30 bins of the trial. Accuracy and confusion matrices were estimated using distinct conditions for training and testing (e.g. in Figure 6H) or five-fold (Figure 6I) or ten-fold cross validation (other figures).

## Supplementary Figures

**Supplementary Figure 1.**
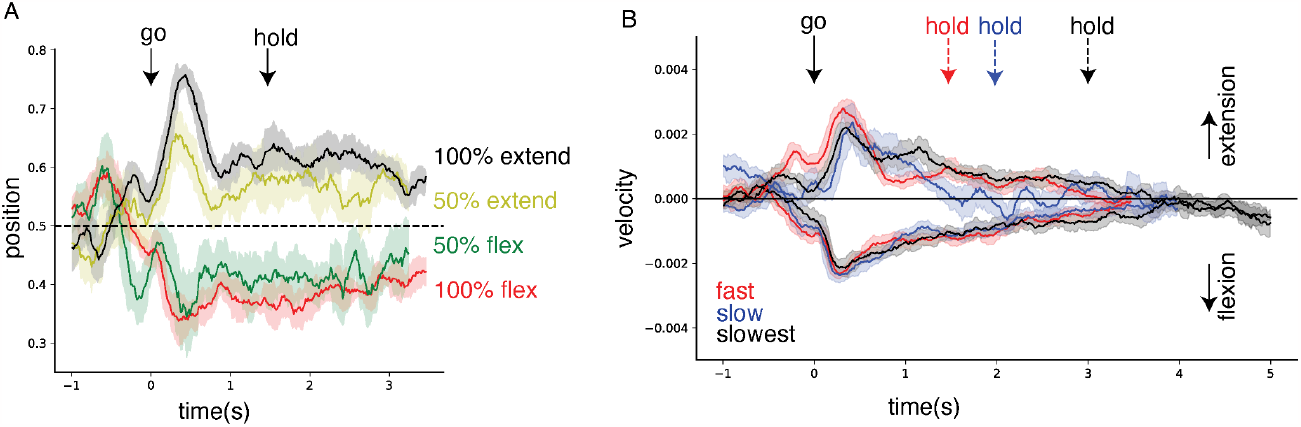
Finger position and velocity decoded from neural activity. (A) T5 attempted thumb movements for different degrees of flexion/extension, and at varying speeds. Predictions of a linear position decoder for a fixed speed and varying degrees of flexion and extension during the hold period (different colors). Start of go and hold periods indicated. Positions along 0-1, with 0.5 being rest, 0 being 100% flexion and 1 being 100% extension. (B) Same as (A), but for velocity decoding.

**Supplementary Figure 2:**
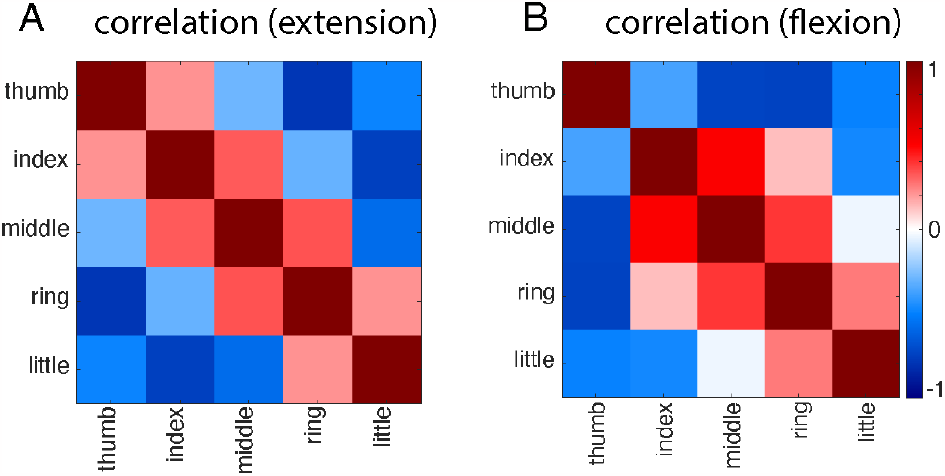
(A) Angular geometry of all fingers during extension is estimated by measuring the Pearson correlation between all the movement conditions. (B) Similar to (A), for flexion.

**Supplementary Figure 3:**
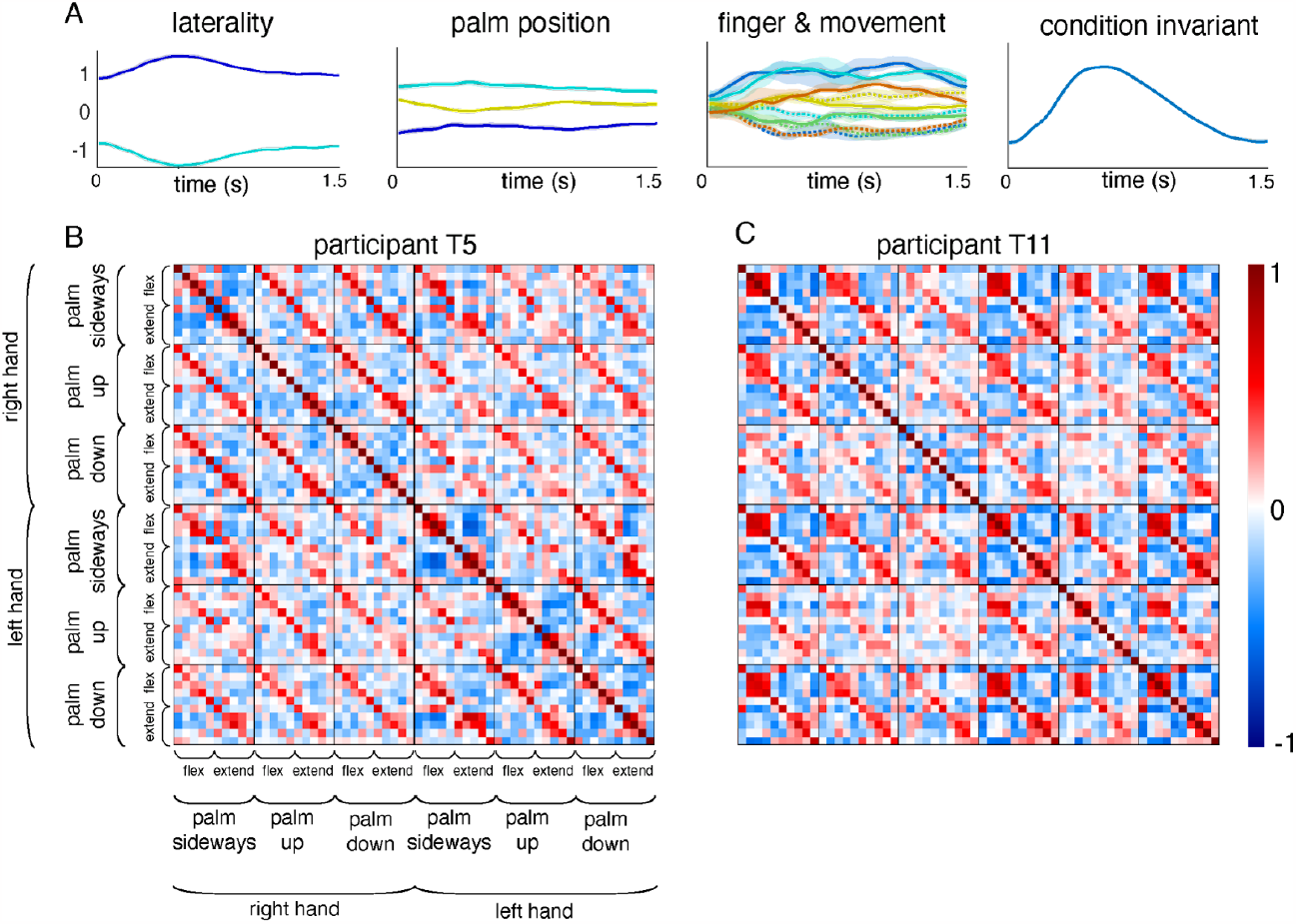
Compositional representation of finger movements across palm position and limb laterality. (A) Top principal component (PC) of the temporal contribution of individual factors (hand laterality, palm position and effector-movement pair) in T5’s neural activity. First, the neural activity is averaged over all trials to get a condition invariant signal and the projection along the first PC is plotted. Second for each factor, the condition invariant activity is subtracted, averaged across all trials for each level of the factor, followed by projection along the top PC. The PC projection for each level of laterality (left hand & right hand), palm position (palm up, down & sideways) and effector (each finger & all fingers together)-movement (flexion & extension) combination is plotted for the 1.5s of trial following the Go cue. (B) Correlation of the time and trial averaged neural activity (from participant T5, multi-unit threshold crossings) between different combinations of hand laterality, palm position, effector and movement type. First, the translational component is removed by subtracting the average activity across all conditions with same hand laterality and palm position, Next, correlation between the resulting neural activity vectors (with length equal to the number of channels) is computed (see Methods). Hence, each smaller square along the diagonal captures the relative geometry within the corresponding context - defined by a particular hand laterality and palm position. Other small squares indicate the similarity of geometry between contexts after removing the context-dependent translation. The observed off-diagonal bands suggest that the relative organization is preserved across contexts. (C) Similar to (B), using local field potentials in participant T11.

**Supplementary Figure 4:**
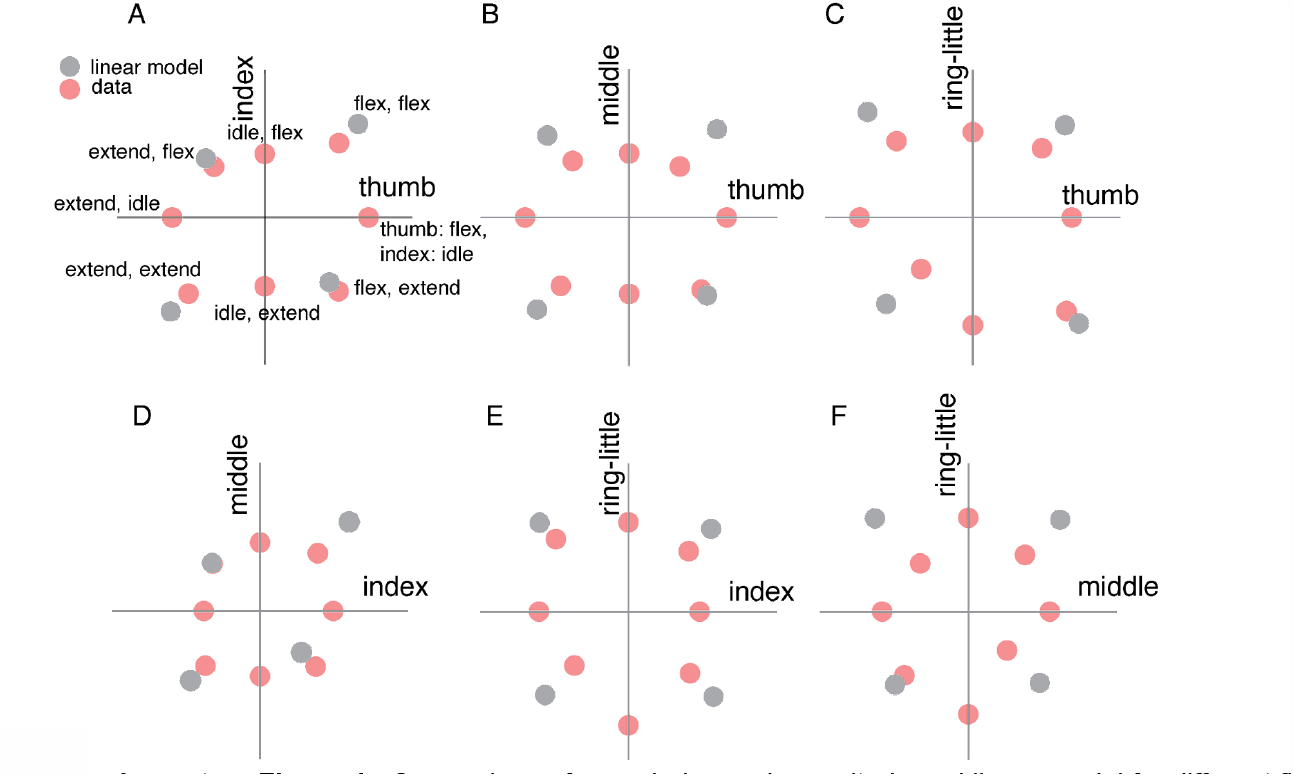
Comparison of recorded neural magnitude and linear model for different finger combinations. Similar to Figure 5A. Neural activity preprocessed by removing condition invariant signal. Each point denotes a unique condition, with the direction corresponding to the flexion/extension/rest for the two finger groups (indicated by x-axis and y-axis labels) and the length corresponding to an unbiased estimate of the magnitude of neural activity. Red dots correspond to the trial averaged neural activity, concatenated across five sessions and the black dots correspond to the predicted activity for simultaneous movements from the summation of corresponding individual movements.

**Supplementary Figure 5:**
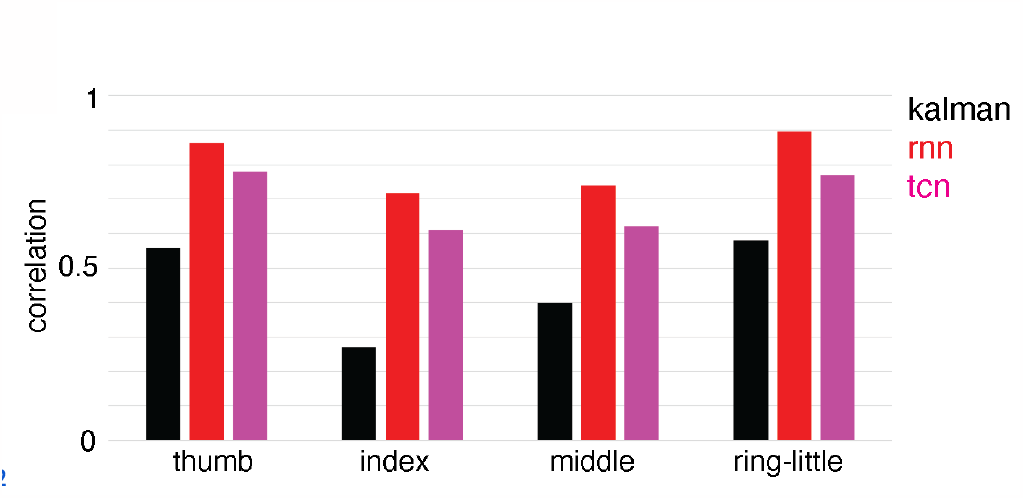
Accuracy of predicting finger positions for open-loop data collected for combinatorial four-finger group movements. Accuracy measured as correlation between cued and predicted trajectory for each finger group in the held-out data. Models include Kalman Filter (black), recurrent neural network (rnn, red) and temporal convolutional network (tcn, magenta). RNN and TCN models were trained using various data augmentation methods (scaling, stretching and additive noise) and hyper-parameters were chosen based on a separate validation dataset. Linear model (Kalman filter) performs worse for index and middle fingers, and the non-linear decoders (rnn and tcn) show a greater improvement for these fingers.

**Supplementary Figure 6:**
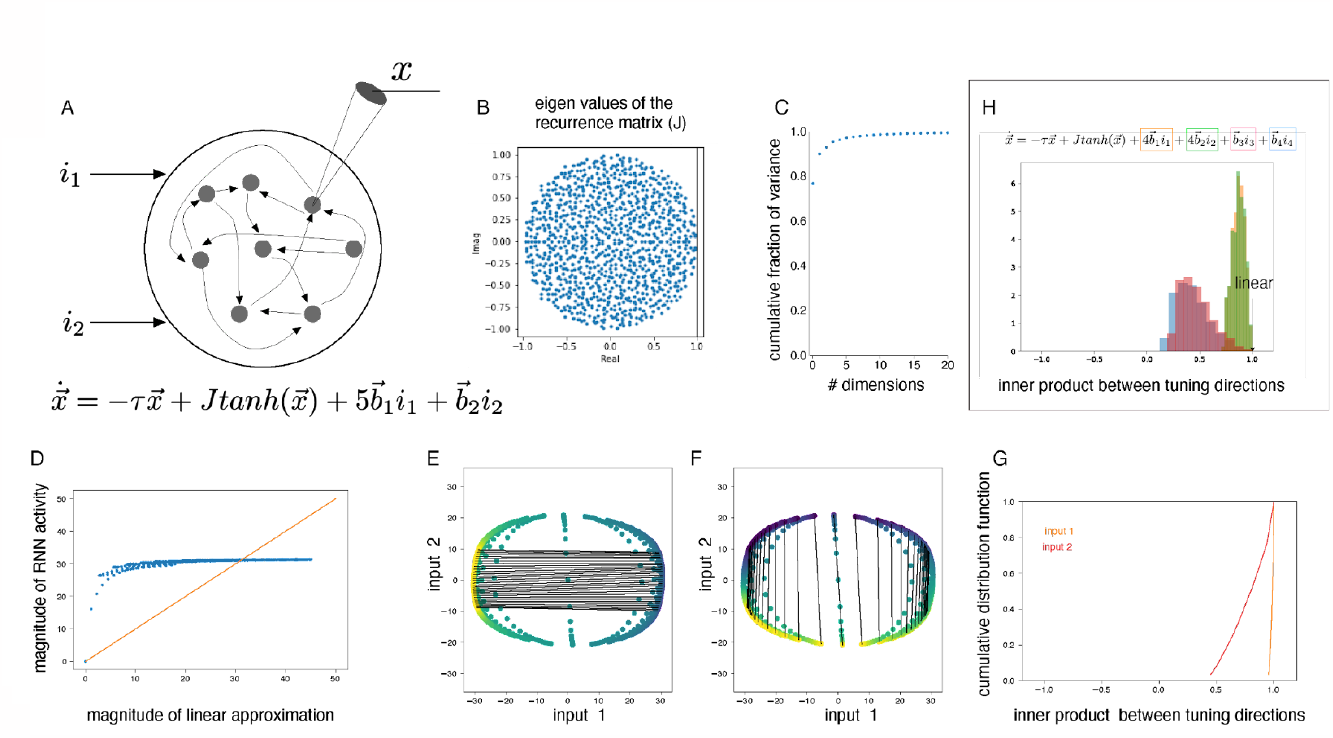
Linear inputs transformed by non-linear RNN captures observed properties. (A) Model schematic. Two scalar inputs (i_1_, i_2_) are processed by a RNN consisting of random connectivity (J) and tanh activation non-linearity. Input 1 is five times stronger than input 2. Different values of static inputs (i_1_, i_2_) are presented and the RNN is simulated till steady state. See Methods for details. Model simulated with 1000 neurons. (B) Eigenvalues of the random connectivity matrix J, uniformly distributed on the complex disk. (C) Dimensionality of the steady-state neural activity for all combinations ranging from -10 to 10 for each input independently. The simulated model captures the low dimensionality of the observed data. (D) The magnitude of RNN simulated neural activity (y-axis) is constant for most inputs and a linear model fails to capture this magnitude invariance. The non-linear RNN captures the normalization effect presented in Figure 5. (E) Population tuning direction of input 1 indicated by lines joining points corresponding to extreme values of input 1, while keeping input 2 fixed. Different lines correspond to different fixed values of input 2. (F) Similar to (E), but population tuning directions for input 2. (G) Greater change in population tuning direction observed for the (weaker) input 2, as indicated by distribution of angles between population tuning directions for both fingers. (H) Extension of the RNN model to four inputs, and repeating analysis from Fig. 6G.

**Supplementary Movie 1.** Animation of Figure 6F, rotated and viewed from different directions. Each finger group is presented sequentially.

